# Habitat and seasonal drivers of leukocyte profiles within and across Neotropical bat species

**DOI:** 10.1101/2025.07.23.666480

**Authors:** Daniel J. Becker, Kristin E. Dyer, Lauren R. Lock, M. Brock Fenton, Nancy B. Simmons

## Abstract

Land conversion is a widespread form of environmental change that can alter infection dynamics in wildlife by modifying host immune defense. Such effects may be compounded by seasonal variation in resources and reproduction and differ among members of a host community, yet the combined effects of habitat, season, and species identity on wildlife immunity remain poorly understood. We tested within- and across-species effects of land conversion and seasonality on immunity in Neotropical bats by quantifying hematological markers of physiological stress and inflammation. We sampled seven species across a large forest preserve and smaller nearby forest fragment in northern Belize during the dry and wet seasons. Using phylogenetic generalized linear mixed models, we tested overall effects of habitat and season and quantified per-species impacts. Total leukocyte counts and neutrophil-to-lymphocyte ratios showed no overall habitat or seasonal effects but had strong species-specific responses to these predictors. In contrast, the systemic inflammation response index was higher across species in the dry season and in the smaller fragment, suggesting poor health in unfavorable conditions. Species-specific effects did not align with diet guilds, indicating potential roles for finer-scale ecological traits. Our results highlight the complex, species-dependent effects of environmental change on wildlife immunity.

## Introduction

Land conversion is one of the dominant and accelerating forms of contemporary environmental change [1], with cascading effects on wildlife population and community dynamics [2,3]. Such changes also have driven increased infectious disease risks within wildlife hosts and, through zoonotic spillover, to humans [4–6]. One of the primary proposed mechanisms by which land conversion alters infection dynamics is through impacts on wild host immunity [7,8]. Decreased amount or quality of space and food resources in degraded or fragmented habitats can increase physiological stress, such as by expending more energy during foraging or increasing frequency of negative intra-or interspecific interactions (e.g., crowding or edge effects) [9,10]. Prolonged physiological stress (i.e., allostatic overload) can in turn affect host immune responses [11,12], including increased susceptibility to new infections and the relapse of chronic infections [13,14].

The immunological consequences of land conversion could be further compounded by seasonality [8]. Seasonal reproductive activity can favor allocation of energetic resources away from immune defenses in mammals, particularly when females are pregnant or lactating [15,16]. Similarly, periods with less-favored climatic conditions or reduced food availability, such as winters in temperate regions or the dry season in tropical regions, can also impose energetic costs and limit energy allocation to immunity [17–19]. As such, habitat degradation or fragmentation could have the most pronounced effects on host defenses during birth pulses or these unfavorable periods, given further increases to energy required for homeostasis and amplifying tradeoffs between immunity and other physiological functions [20,21]. For example, such interactive effects were hypothesized to explain why pulses of Hendra virus shedding from Australian flying foxes occur primarily in the winter months and for colonies that recently experienced food shortages and were displaced into agricultural and urban habitats [22]. However, the immunological underpinnings of such relationships remain poorly understood, partly because their study requires field sampling designs that jointly capture the effects of both habitat and seasonality [23]. More even host sampling across habitats and among seasons could test hypotheses about interactive effects and reveal where and when hosts are susceptible [24].

Lastly, although first principles surrounding energy expenditure and allocation suggest overall deleterious impacts of land conversion and unfavorable seasons on wildlife immunity, this outcome may not be homogeneous across host species. As emphasized through trait-based approaches to studies of global change [25], species do not respond equally to increasing anthropogenic pressures, even if mean effects are often negative [26–28]. For example, the demography and distribution of some generalists are less affected by habitat fragmentation than specialists [29,30], suggesting such species could show less immunological change in converted habitats. Many generalists can capitalize on new resources provided in these altered landscapes, potentially allowing greater allocation of energy into immunity [31,32]. For example, white ibis occupying urban habitats feed more consistently on anthropogenic resources and, in turn, have lower glucocorticoid levels and stronger innate immunity [33,34]. From a similar perspective, species that rely on more ephemeral resources, such as frugivores or nectarivores, may be especially physiologically vulnerable to seasonal shifts in food resources [35], while species with generalist foraging behavior could instead show less seasonal variation in immunity [36]. Given such complexities, there is a need for spatially and temporally explicit field studies that assess immune impacts within not only a single host species but also across the broader community.

We here provide an initial test of the within- and across-species effects of land conversion and seasonality on immunity in the context of Neotropical bats. Bats (order Chiroptera) have been increasingly studied as model systems for comparative immunology [37], given multiple immune adaptations that differ from other mammals and substantial inter-specific variation in defense [38,39]. Bats display species-specific responses to land conversion [40,41], with especially pronounced effects in the Neotropics [42]. The Neotropics contain some of the most extreme concentrations of bat diversity globally [43], driven in part by diversification through dietary shifts [44,45]. Neotropical bats encompass a wide range of foraging strategies, including not only frugivory, nectarivory, and insectivory but also carnivory, piscivory, and hematophagy [46,47]. Land conversion for cropland and pasture is also widespread and accelerating in the Neotropics [48,49], with the responses of bat demography and occupancy dependent on foraging ecology [50,51]. Diet also plays a role in immune differences among Neotropical bat species, their physiological responses to resource availability, and if they show unimodal or bimodal reproductive phenologies [52–56]. Neotropical bats are thus an ideal system for testing species-specific responses of host immunity to land conversion and season.

We focused our analyses on a long-term research program in northern Belize (Orange Walk District), where increasing habitat fragmentation driven by agricultural change has led to pronounced shifts in bat communities [57–59]. Surveys at the Lamanai Archeological Reserve (LAR) and Ka’Kabish (KK), which comprise a large preserve and a small, isolated fragment, respectively, indicate the KK bat community is a nested subset of that found in the LAR [59]. Our prior work here has shown site-level differences in contaminant dynamics (in part driven by agricultural change) that impact bat immunity [60] as well as select species-specific responses of bat immunity to ongoing habitat fragmentation in the region for bats occurring in the LAR [61]. Other efforts have focused on immunological characterization of those species that occur in both sites, such as *Desmodus rotundus, Artibeus jamaicensis*, and *Pteronotus mesoamericanus* [62– 65]. By systematically sampling common bat species in both the LAR and KK and across two seasons, we test the hypothesis that bat immune systems, as revealed by hematology, would show signatures of physiological stress and inflammation that are most pronounced in the smaller habitat and unfavorable season. At the same time, we predicted that bat species would also vary in their immunological response to both habitat and season in ways that align with foraging ecology, such as by those that can capitalize on year-round prey in agricultural habitats (e.g., insectivorous and hematophagous bats) being less physiologically affected by agriculture and seasonality, while frugivorous and nectarivorous species dependant on forest plants could be especially immune impaired in unfavorable environments owing to reduced food availability.

## Methods

### Bat sampling

During two-week periods in November 2021 and April–May 2022, we sampled bats in both the LAR and KK as part of broader studies of intra- and interspecific variation in immunity and pathogen diversity [60,61,66]. These periods coincided with the late wet season (November) and the height of the dry season (April–May) [67], with climatic differences being driven mostly by temperature rather than by precipitation (Fig. S1). Bats were captured using mist nets and harp traps along flight paths, held in individual bags, and identified to species based on morphology [59,68]. We also recorded sex, reproductive status, and age [69]. We collected blood using heparinized capillary tubes after lancing the propatagial vein with sterile needles (26–30G); all blood volumes were less than 1% body mass. Thin blood smears were prepared on glass slides, air dried in the field, and stained with Wright–Giemsa (Astral Diagnostics Inc., Hematology Stain Set Quick III) at the University of Oklahoma. All bats for this study were released following sampling. Field procedures were performed according to guidelines for the safe and humane handling of bats published by the American Society of Mammalogists [70] and were approved by the Institutional Animal Care and Use Committees of the American Museum of Natural History (AMNH IACUC-20210614) and University of Oklahoma (2022–0197). All bat sampling was authorized by the Belize Forest Department through scientific collection permit FD/WL/1/21(12) and Belize Institute of Archaeology permits IA/S/5/6/2l(01) and IA/H/1/22(03).

### Hematological analyses

Using blood smears, we first estimated total white blood cell (WBC) counts as the mean number of leukocytes from 10 fields of view (400X) with a light microscope [54]. We next used differential WBC counts (1000X, oil immersion) to quantify the relative abundance of neutrophils, lymphocytes, monocytes, eosinophils, and basophils from 100 leukocytes [71]. We used our differential WBC counts to derive hematological indices of physiological stress and inflammation, including the neutrophil-to-lymphocyte ratio (NLR) and systemic inflammation response index (SIRI, a ratio of the neutrophil and monocyte quotient to lymphocytes) [72–74].

### Statistical analyses

From a broader sample of 294 bats (*n* = 21 species) with hematology data, we focused analyses on species captured in both the LAR and KK and with a sample size greater than or equal to nine (*n* = 232 individuals spanning seven species). These seven species included *Desmodus rotundus* (*n* = 77), *Artibeus jamaicensis* (*n* = 48), *Sturnira parvidens* (*n* = 29), *Pteronotus mesoamericanus* (*n* = 25), *Glossophaga mutica* (*n* = 23), *Carollia sowelli* (*n* = 21), and *Saccopteryx bilineata* (*n* = 9), representing one hematophagous species (*D. rotundus*), three frugivorous species (*A. jamaicensis, S. parvidens, C. sowelli*), one nectarivorous species (*G. mutica*), and two insectivorous species (*P. mesoamericanus* and *S. bilineata*). All species belong to the family Phyllostomidae with the exceptions of *P. mesoamericanus* (Mormoopidae) and *S. bilineata* (Emballonuridae; Fig. S2). Per-species sample sizes varied among seasons and sites (Fig. S3), and species varied in the seasonal distribution of male and female reproductive activity (Fig. S4). Within this dataset, our hematology responses were not correlated (*ρ* = –0.14 to 0.54, 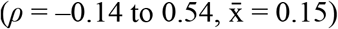).

We used phylogenetic generalized linear mixed models (PGLMMs) fit using the *brms* package in R to test effects of site and season on hematology while controlling for evolutionary history [75,76]. For each of our hematological outcomes (total WBC count, NLR, SIRI), we included the above fixed effects alongside sex, reproductive status, and age. We also included the time between capture and blood collection (i.e., holding time) to account for impacts of this acute stressor [72,77]. Because a small number of bats lacked data on reproductive status (*n* = 5), age (*n* = 5), or holding time (*n* = 9), we used the *missRanger* package to impute these missing values using chained random forests (*n* = 1,000 trees) [78,79]. We assessed collinearity by fitting a linear model with the same fixed effects, finding consistently low variance inflation factors (1.04–1.48). To account for species-specific responses, each model included a random intercept of species and random slopes of site and season [80]. We also included a phylogenetic random intercept using a variance–covariance matrix derived from the bat phylogeny using the *ape* package [81,82]. We did not estimate the correlation between random slopes and intercepts nor included phylogenetic random slopes to avoid overparameterization and to improve PGLMM convergence [83]. All models used Gaussian errors with log_10_-transformed response; owing to zero values for the total WBC count and SIRI, we added half the minimum non-zero value prior to transformation [84]. We used leave-one-out information criterion (LOOIC) to compare our base model to a model that also included the interaction between site and season [85]. We ran our models using four chains for 12,500 iterations and a burn-in of 50%, thinned every 25 steps. We verified model convergence by inspecting trace plots and 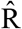 values. We then summarized the posterior mean and corresponding 95% credible interval per each PGLMM coefficient, including species-specific slopes for both site and season, using the *ggdist* and *tidybayes* packages [86]. Lastly, we also assessed both marginal and conditional *R*^*2*^ with the *performance* package [87,88].

## Results

Across all three hematology outcomes in our sample subset of the Belize bat community, PGLMMs with additive effects of site and season were always supported by LOOIC over models with interactive effects (Table S1). The fixed effects explained 10–22% of the variation in our hematology variables, while the species-level random effects explained an additional 5–15% of the variance. For total WBC counts and the NLR, we found no overall effect of site or season (Table 1, Fig. 1). However, males had higher total WBC counts, and reproductively active bats had lower total WBC counts and elevated NLRs. Our models found no baseline difference in the total WBC count nor species-specific responses to seasonality (i.e., low variation in the species-level random intercepts and in the random slopes of season), whereas we did find species-specific responses to site (i.e., non-zero random slopes of habitat). We also detected baseline differences in the NLR and species-specific responses to season (i.e., non-zero random intercepts of species and random slopes of season) but no species-specific responses to site (i.e., low variation in the random slopes of site). By contrast, the SIRI did show overall effects of both site and season, with this measure of inflammation being elevated in KK and the dry season (Table 1, Fig. 1). Bats also displayed species-level differences in baseline SIRI measures (i.e., a non-zero random intercept of species), whereas the SIRI did not show species-level variation by site or season (i.e., low variation in random slopes). None of our models found strong age or residual phylogenetic effects, and only the NLR showed a positive association with holding time.

**Table 1.** Posterior means and 95% credible intervals (highest density intervals) of fixed and random effects from the top PGLMMs for all three hematological response variables in Belize bats. Parameters for which 95% credible intervals do not cross zero are displayed in bold.

| Effects | Parameter | total WBC | NLR | SIRI |
| --- | --- | --- | --- | --- |
| Fixed<br>($\beta$) | Intercept | -0.28 (-0.75, 0.37) | **-0.47 (-0.96, -0.03)** | **-2.36 (-3.03, -1.56)** |
|  | KK | 0.02 (-0.28, 0.34) | 0.12 (-0.14, 0.42) | **0.44 (0.03, 0.86)** |
|  | Dry season | -0.11 (-0.31, 0.08) | 0.13 (-0.21, 0.45) | **0.50 (0.15, 0.83)** |
|  | Males | **0.20 (0.09, 0.32)** | -0.01 (-0.14, 0.13) | -0.11 (-0.37, 0.13) |
|  | Adult | -0.04 (-0.20, 0.16) | 0.16 (-0.02, 0.37) | 0.03 (-0.35, 0.40) |
|  | Reproductive | **-0.16 (-0.31, -0.01)** | **0.35 (0.20, 0.52)** | 0.27 (-0.06, 0.58) |
|  | Holding time | 0.06 (-0.20, 0.28) | **0.28 (0.04, 0.60)** | 0.16 (-0.37, 0.67) |
| Random<br>(SD) | 1\|species | 0.209 (0, 0.519) | **0.176 (0.001, 0.429)** | **0.248 (0.001, 0.637)** |
|  | site\|species | **0.301 (0.004, 0.687)** | 0.217 (0, 0.559) | 0.254 (0, 0.698) |
|  | season\|species | 0.151 (0, 0.377) | **0.324 (0.005, 0.721)** | 0.216 (0, 0.55) |
|  | 1\|phylogeny | 0.048 (0, 0.109) | 0.035 (0, 0.083) | 0.052 (0, 0.119) |

**Figure 1.**
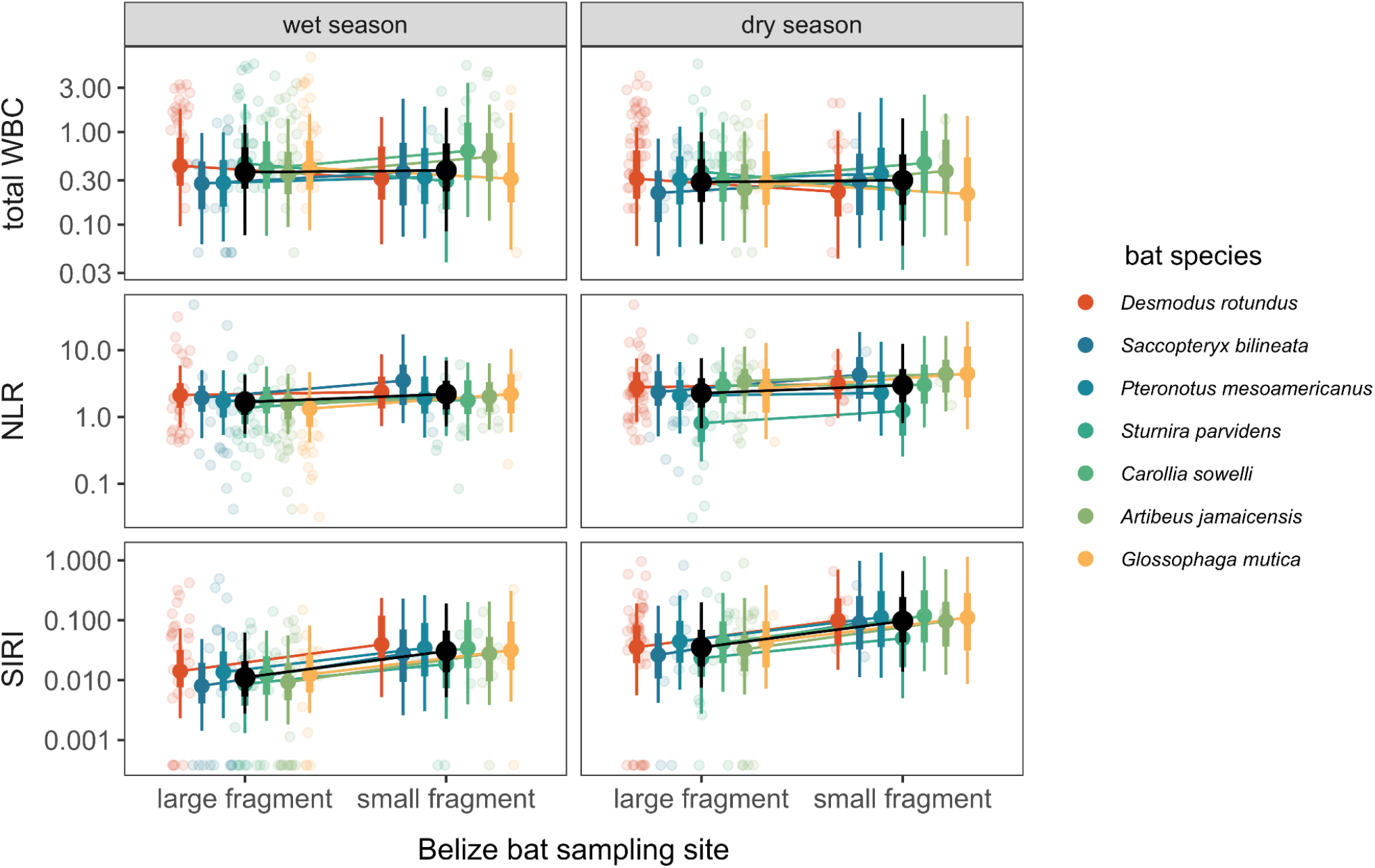
Posterior means and credible intervals (highest density intervals, 66% and 95%) are shown for the additive effects of habitat and season (black) for all three hematology PGLMMs. Species-specific responses derived from each model’s random effects are also shown, with raw data overlaid and jittered to reduce overlap. Hematology values are displayed on a log_10_ scale.

Although bats showed species-specific hematological responses to season (i.e., NLR) and site (i.e., total WBC counts), these effects did not map onto dietary guild (Fig. 2). Most species had fewer WBCs in the dry season, with the exception of *Pteronotus mesoamericanus*; however, 95% credible intervals for all dry season effects crossed zero. By contrast, most species had elevated NLRs in the dry season, with this effect strongest for *Artibeus jamaicensis*; only *Sturnira parvidens* had lower NLRs during this period. Given the negligible random slopes of season for SIRI and strong overall effect, all bats had elevated inflammation in the dry season, although effects were strongest for *Pteronotus mesoamericanus, Carollia sowelli*, and *Artibeus jamaicensis*. Occupying KK had variable species-specific effects for the total WBC count, with both insectivorous species, *Carollia sowelli*, and *Artibeus jamaicensis* having more leukocytes while *Desmodus rotundus, Sturnira parvidens*, and *Glossophaga mutica* had fewer leukocytes (although 95% credible intervals for these KK effects crossed zero). Most bats had weakly higher NLRs in KK, with effects strongest for *Saccopteryx bilineata* and *Glossophaga mutica* (all 95% credible intervals again crossed zero). Given the overall effect of site on the SIRI, all species had elevated inflammation in KK, but this contrast was greatest for *Desmodus rotundus* and *Artibeus jamaicensis* (whereas 95% credible intervals crossed zero for all other species).

**Figure 2.**
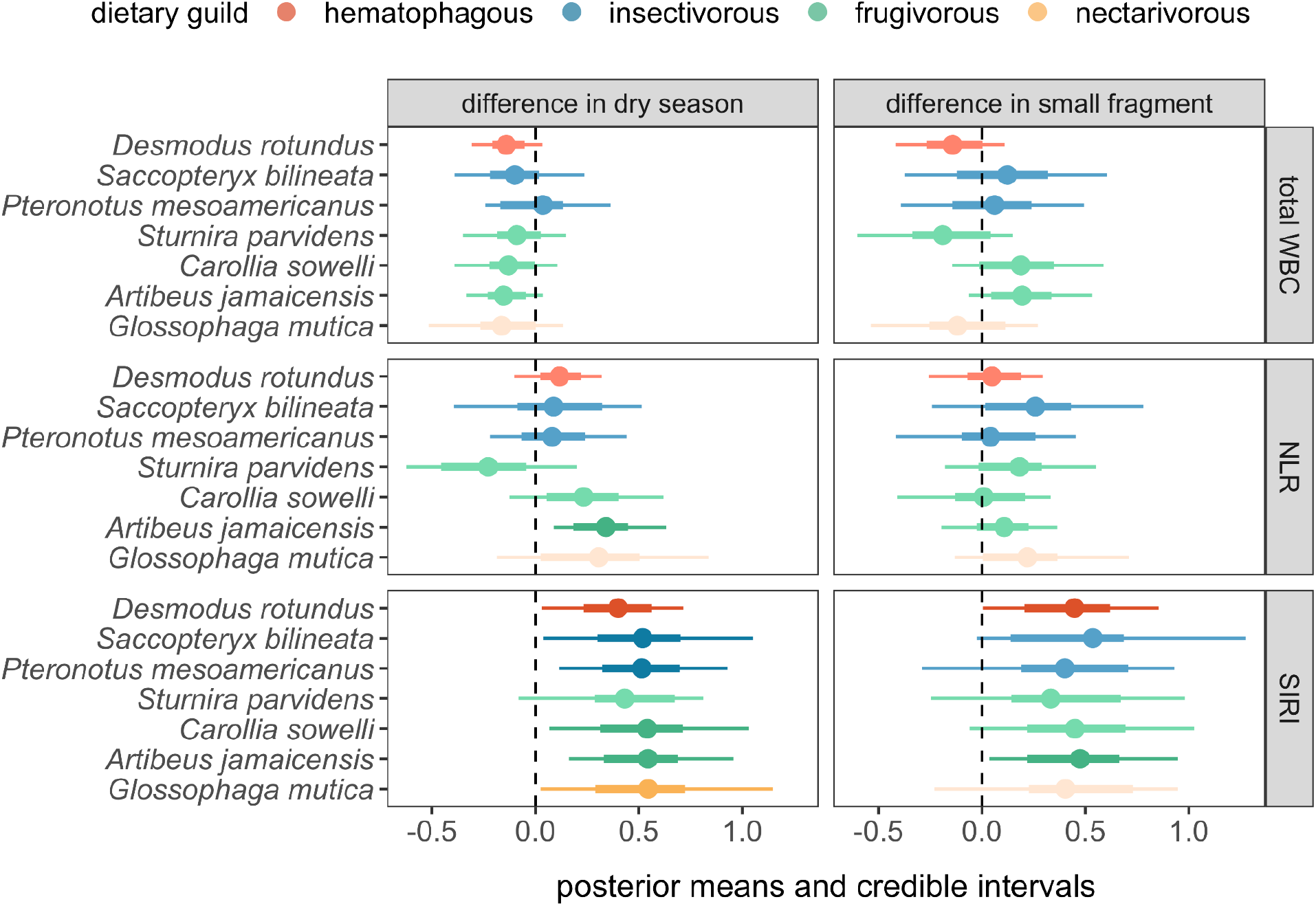
Posterior means and credible intervals (highest density intervals, 66% and 95%) are shown for the species-specific habitat and season coefficients derived from each hematology PGLMM. Effects are colored by dietary guild, with darker shading indicating those effects in which the 95% credible intervals do not overlap with zero (dashed vertical line). Estimates are presented after statistically adjusting for sex, reproductive status, age, and log_10_-holding time.

## Discussion

Although land conversion is hypothesized to affect infectious disease risks through changes in wild host allostatic load and, in turn, immune defense [7,8,10,21], how these effects are modified by seasonal variation in wildlife energetics as well as by host ecology and evolution remain poorly understood. Here, we leveraged a long-term study system on diverse Neotropical bats in Belize to partition the within- and across-species effects of habitat fragmentation and seasonality on cellular immunity. While some hematological indices did display consistent effects of site and season (e.g., the SIRI), other measures showed stronger species-specific responses to these drivers. However, these species-specific effects did not align clearly with dietary guild, indicating roles for finer-scale traits describing foraging ecology or other characteristics such as reproductive phenology or roosting ecology. Together, our findings demonstrate the complex effects of land conversion on wildlife immune defense and highlight important next steps to improve our understanding of how environmental change ultimately affects infection risks.

Our work provides general support for plausible immune impairment in fragmented habitats and the dry season. Specifically, the SIRI was elevated in both contexts across the bat community (i.e., 95% credible intervals for the fixed effects did not include zero). The SIRI is a biomarker of inflammation originally validated in human cancer studies, involving neutrophilia, lymphopenia, and monocytosis as well as increased levels of some inflammatory cytokines [73]. A relative increase in neutrophils and decrease in lymphocytes in blood (i.e., increased NLR) often indicates prolonged physiological stress, whereas increases in the relative abundance of monocytes are often interpreted to indicate acute infection [72]. We observed no overall increase in the NLR in the smaller fragment or dry season, although work in captive vertebrates suggests monocytosis can also be a result of stressors including but not limited to heat stress, crowding, and pregnancy [89–92]. While we cannot discount the possibility that elevated infection risks, especially with known bacterial and protozoan pathogens common in this study system [60,62], are likewise a cause of observed habitat and seasonal differences in the SIRI, such drivers could also result from feedbacks between poor condition and infection [93]. More broadly, however, our findings are consistent with the hypothesis that energetic tradeoffs and allostatic load can be exacerbated by land conversion and unfavorable periods, resulting in inflammation and immune dysregulation [7,8,21]. Our overall seasonal results may also be driven in part by reproduction, as general reproductive activity was associated with lower total WBC count, higher NLR, and weakly higher SIRI across bat species; furthermore, most reproductive activity across male and female bats was observed in the dry season (69%) compared to the wet season (41%; Fig. S4). Such findings are consistent with tradeoffs between reproduction and immunity observed in multiple wildlife systems, including but not limited to bats [15,16,94–96], although direct effects of climate and indirect effects of resource limitation during the dry season cannot be discounted.

While we found overall effects of site and season on the SIRI, we observed stronger heterogeneity among species in how the total WBC count and NLR responded to these stressors. For example, *Artibeus jamaicensis* had the strongest seasonal increase in the NLR compared to other species, with some species such as *Sturnira parvidens* showing a weak decrease in the dry season. Frugivorous species also differed in habitat effects on total WBC counts. Similarly, both *Desmodus rotundus* and *Artibeus jamaicensis* had elevated SIRI in the smaller fragment, despite having highly contrasting diets. Such examples illustrate how the hematological response of species to unfavorable conditions does not map clearly onto dietary guilds, even though foraging ecology plays an important role in diverse aspects of bat biology, including but not limited to demographic responses to land conversion and immunological variation [50–56,61]. This finding underscores caveats about traditional guild assignments, as species may not fit neatly into such categories [46,47] and species-level differences in the immunological response to environmental conditions may be driven by other foraging factors such as degree of dietary specialization [97].

Future work is needed to better understand the characteristics that predispose species to have different immunological outcomes in response to land conversion and seasonality [25,98]. Reproductive phenology may be one such candidate trait, given some concordance between the seasonal distribution of reproductive activity in our data and species-specific effects of the dry season on cellular immunity. For example, for *Artibeus jamaicensis*, for which we found elevated NLRs in the dry season, we only observed pregnant females in this same season, and lactation was more common in this period than in the wet season (Fig. S4); this reproductive phenology is similar to that found elsewhere in the species range [99]. Similarly, we only found reproductive females in the dry season for *Pteronotus mesoamericanus* and *Carollia sowelli* (Fig. S4), again consistent with reproductive phenologies for these species or their related counterparts elsewhere in Central America [100], with both species also showing strong increases in the SIRI during the dry season. Roosting ecology could be another relevant trait-based predictor, as less-ephemeral roosting sites (e.g., caves and tunnels as opposed to vegetation) could decrease vulnerability to environmental stressors [54,101]. Similarly, body mass could play a role in differentiating these immune responses, given the potential for allometric scaling in immune defense [102,103]; further, some of our species with the strongest response to seasonality and habitat were relatively larger. Ultimately, however, more comprehensive sampling across host species will be needed to facilitate a trait-based understanding of immune responses to such environmental conditions.

Although our work identifies community-wide and species-specific immunological responses to land conversion and seasonality, we did not find support for interactive effects of these spatial and temporal factors. Such effects remain plausible, however, given underlying principles of energetic tradeoffs and the evidence here for both factors influencing bat allostatic load [20,21]. Accordingly, we caution that the lack of support for interactions between habitat and season may be due in part to sample size, as we were unable to evenly sample each species in both sites during both sampling periods due to logistical constraints and other factors such as roost abandonment (Fig. S3). Similarly, whereas our analysis focused on a single year of field data collection, the effects of habitat fragmentation on host immunity and infection can take years to manifest [104], and multi-year sampling would be necessary to differentiate consistent seasonal effects from idiosyncrasies of a single year [23]. As such, future work with more even and long-term host sampling across both space and time will play an important role in robustly testing if outcomes such as inflammation are especially elevated in fragmented habitats during unfavorable periods or reproductive seasons [8,23,24]. Our findings here provide support for the independent effects of these factors that are preconditions for more interactive relationships, and our results more broadly demonstrate the importance of partitioning both within- and across-species effects when considering how environmental change affects wildlife immunity.

## Acknowledgements

We thank Mark Howells, Karen Gonzalez, Neil Duncan, and staff of the Lamanai Field Research Center for assistance with field logistics and permits. We also thank the many colleagues who helped capture bats during 2021 and 2022 bat research trips in Belize.

## Data availability

Individual-level data are available in the Dryad Digital Repository: https://doi.org/10.5061/dryad.d51c5b0g2

