## Supplemental Material for "Habitat and seasonal drivers of leukocyte profiles within and across Neotropical bat species"

Figure S1. Seasonality in temperature and precipitation during the study period
Figure S2. Phylogeny of the included bat species for inter-site and inter-season comparisons
Figure S3. Sample size of included bat species for inter-site and inter-season comparisons
Figure S4. Seasonal distribution of reproductive activity per sex per bat species
Table S1. LOOIC comparison of PGLMMs with and without site-by-season interactions

Figure S1. Points show daily mean precipitation (mm) and temperature (°C) in Orange Walk District, Belize during 2021–2022, with bat sampling periods highlighted by dashed lines. Data were provided by the National Meteorological Service of Belize from Tower Hill, ~30 kilometers from the LAR and KK field sites. Overlaid are the fitted means and 95% credible intervals from generalized additive models fit with restricted maximum likelihood via the *mgcv* R package. Each model included effects of year, a cyclic smooth term for ordinal date, and their interaction. We used a Tweedie response for precipitation and a Gaussian response for temperature.

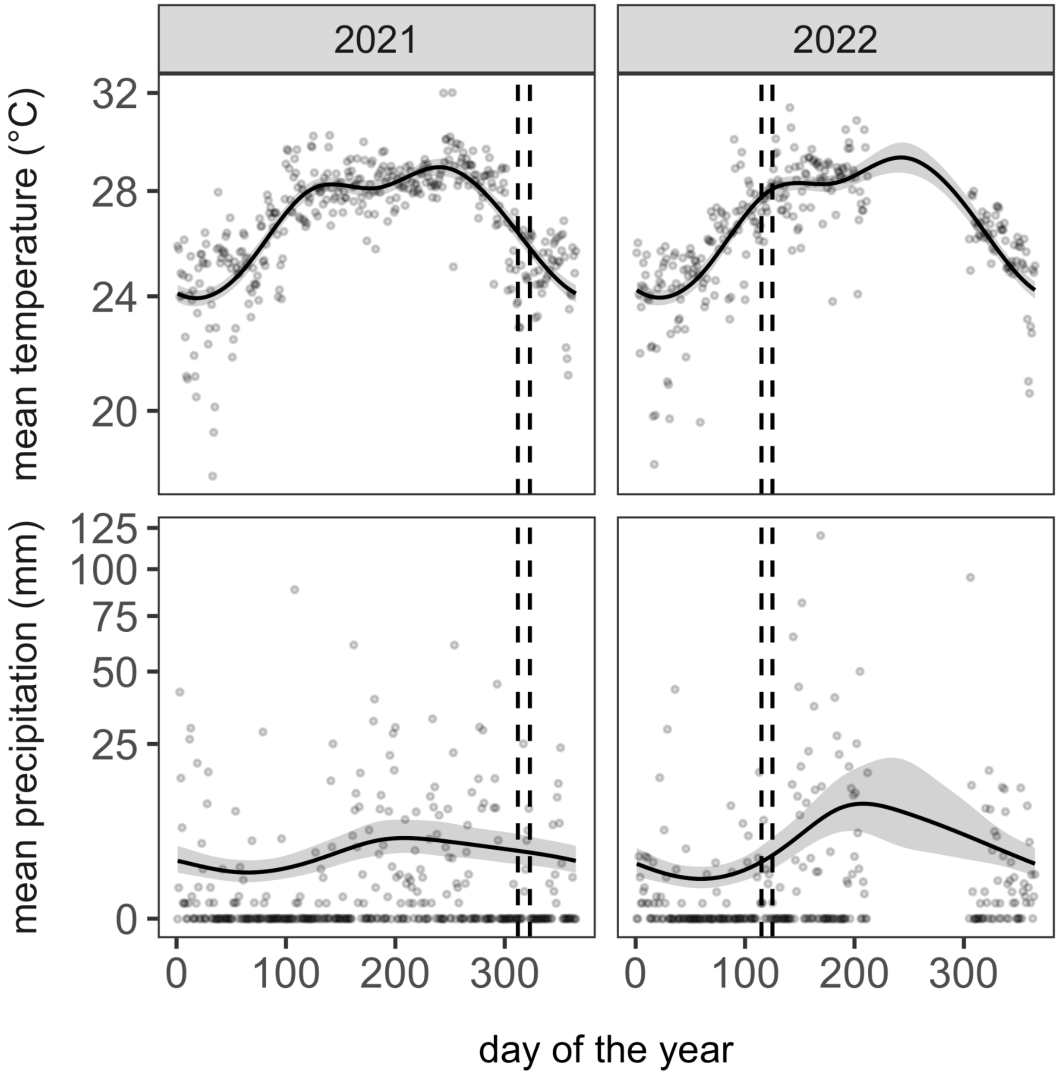

Figure S2. Phylogeny of the included bat species for inter-site and inter-season comparisons.

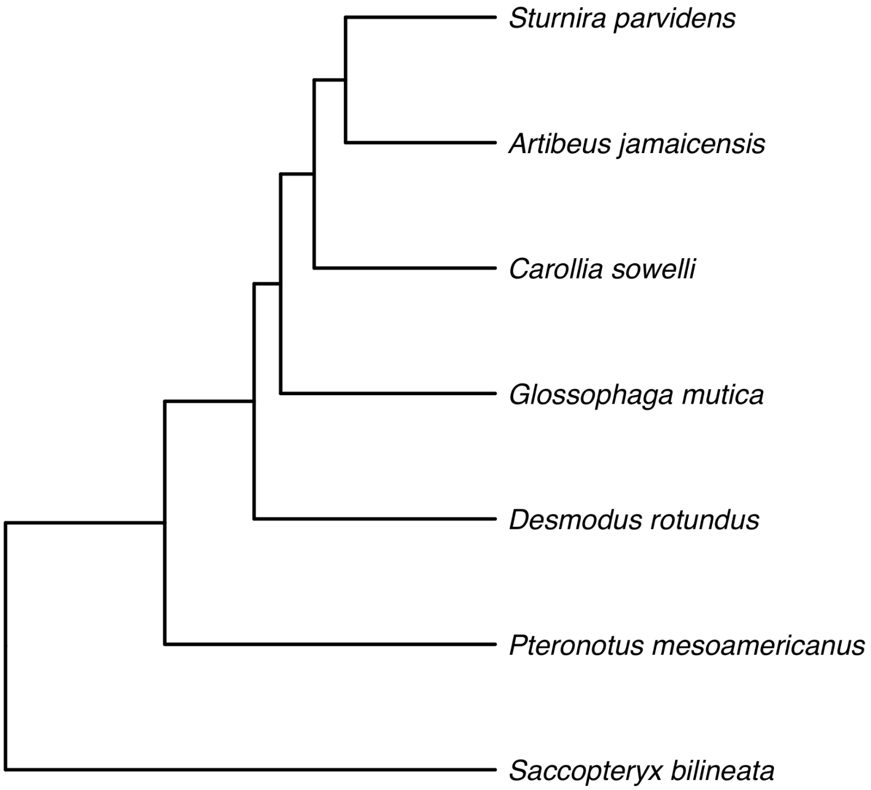

Figure S3. Sample size of included bat species for inter-site and inter-season comparisons.

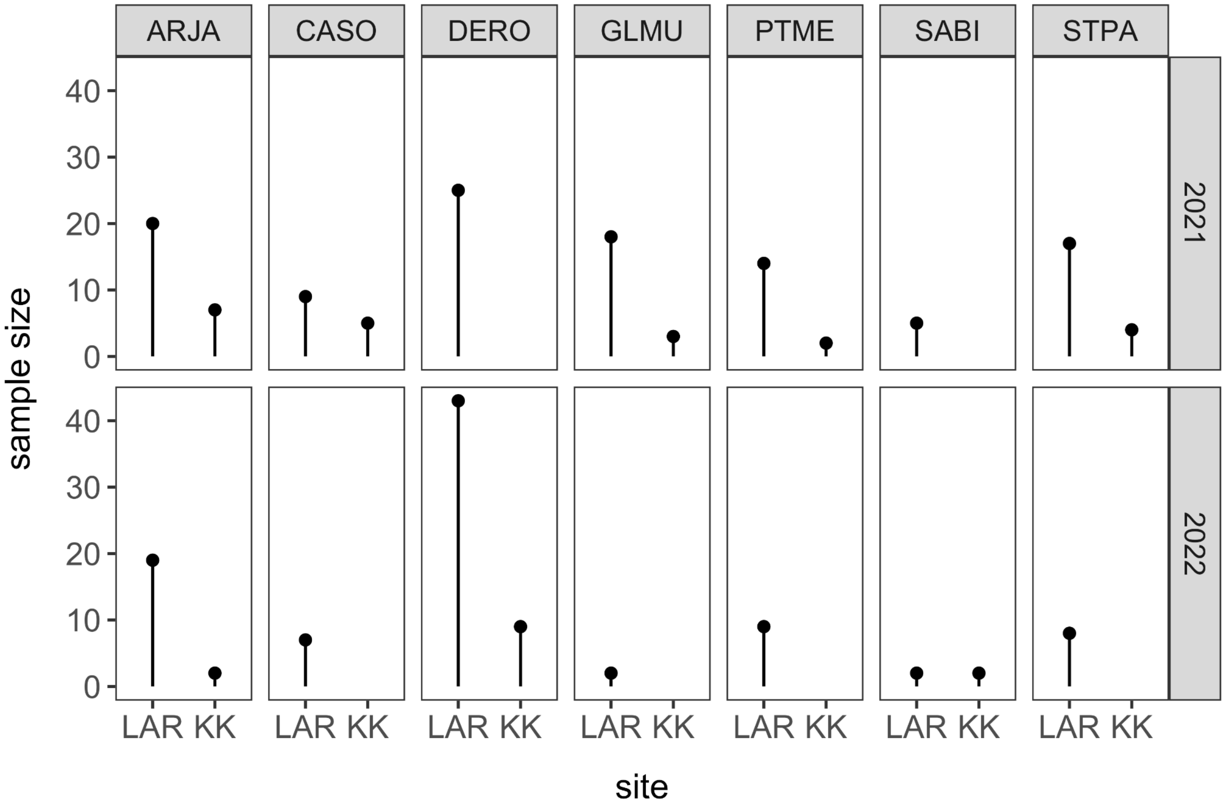

Figure S4. Seasonal distribution of reproductive activity per sex per bat species.

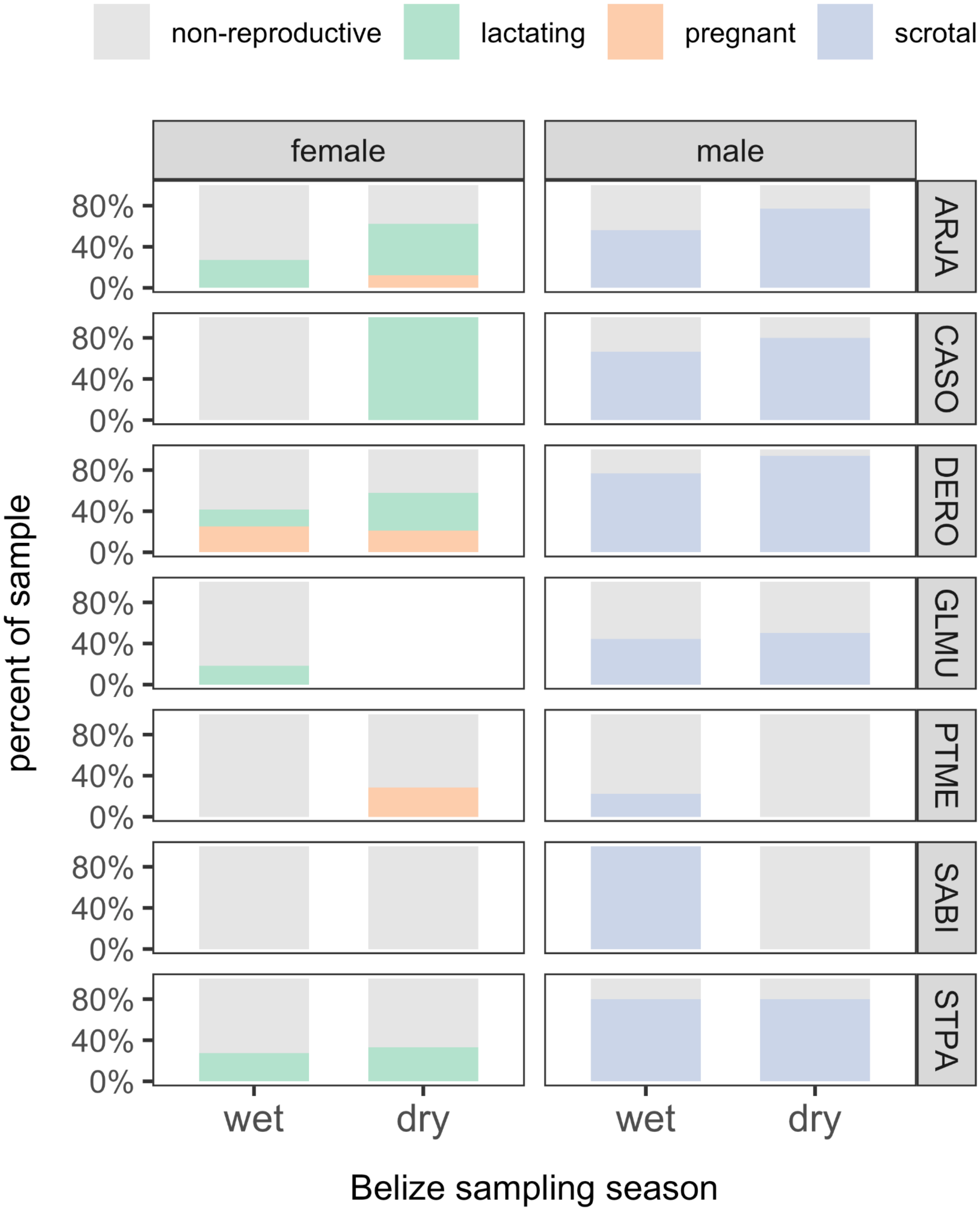

Table S1. LOOIC comparison of PGLMMs with and without an interaction between site and season. All models contained main effects of site, season, sex, age, reproductive status, and $\log_{10}$-holding time as well as a species-level random intercept with random slopes for site and season (as well as a separate phylogenetic random intercept). Models are presented for all three hematology responses with LOOIC (and SE), conditional and marginal $R^2$, and ΔLOOIC.

| **Response** | **Model** | ***k*** | **LOOIC** | **SE** | $R^2_c$ | $R^2_m$ | **ΔLOOIC** |
|---|---|---|---|---|---|---|---|
| Total WBC | All main effects | 7 | 274.05 | 21.88 | 0.25 | 0.10 | 0.00 |
| | + site:season interaction | 8 | 275.54 | 21.83 | 0.26 | 0.11 | 1.49 |
| NLR | All main effects | 7 | 342.27 | 25.65 | 0.35 | 0.21 | 0.00 |
| | + site:season interaction | 8 | 344.02 | 25.70 | 0.36 | 0.22 | 1.75 |
| SIRI | All main effects | 7 | 638.94 | 19.47 | 0.19 | 0.14 | 0.00 |
| | + site:season interaction | 8 | 639.94 | 19.15 | 0.20 | 0.15 | 1.00 |
